# Bio-optical signatures of *insitu* photosymbionts predict bleaching severity prior to thermal stress in the Caribbean coral species *Acropora palmata*

**DOI:** 10.1101/2023.07.06.547971

**Authors:** Kenneth D. Hoadley, Sean Lowry, Audrey McQuagge, Shannon Dalessandri, Grant Lockridge, Eleftherios Karabelas, Courtney Klepac, Carly Kenkel, Erinn M. Muller

## Abstract

The identification of bleaching tolerant traits among individual corals is a major focus for many restoration and conservation initiatives but often relies on large scale or high-throughput experimental manipulations which may not be accessible to many front-line restoration practitioners. Here we evaluate a machine learning technique to generate a predictive model which estimates bleaching severity using non-destructive chlorophyll-a fluorescence photophysiological metrics measured with a low-cost and open access bio-optical tool. First, a four-week long thermal bleaching experiment was performed on 156 genotypes of *Acropora palmata* at a land-based restoration facility. Resulting bleaching responses (percent change in Fv/Fm or Absorbance) significantly differed across the four distinct phenotypes generated via a photophysiology-based dendrogram, indicating strong concordance between fluorescence-based photophysiological metrics and future bleaching severity. Next, these correlations were used to train and then test a Random Forest algorithm-based model using a bootstrap resampling technique. Correlation between predicted and actual bleaching responses in test corals was significant (*p <* 0.0001) and increased with the number of corals used in model training (Peak average R^2^ values of 0.42 and 0.33 for Fv/Fm and absorbance, respectively). Strong concordance between photophysiology-based phenotypes and future bleaching severity may provide a highly scalable means for assessing reef corals.

## Introduction

Increasingly frequent coral bleaching events caused by ocean warming continue to decimate reef systems across the globe (Hughes et al., 2017; Hughes et al., 2018). More than ever before, coral reef conservation initiatives are focused on combating ecosystem loss through the transplant of coral fragments onto impacted sites (Boström-Einarsson et al., 2020; Caruso et al., 2021; Voolstra et al., 2021). Such efforts are meant to mitigate further ecosystem decline while more permanent solutions to ocean warming can be found. The success of many coral restoration initiatives, especially those on heavily impacted sites or areas expected to experience severe environmental perturbations, is reliant on establishing colonies with more environmentally resilient traits (Voolstra et al., 2020; Voolstra et al., 2021; Grummer et al., 2022; Klepac et al., 2023). Indeed, high phenotypic variability in bleaching severity to thermal stress exists within and across individual coral colonies (Parkinson et al., 2015; Kenkel & Matz, 2016), and environments (Kenkel et al., 2013b; Palumbi et al., 2014; Barshis et al., 2018; Voolstra et al., 2020) and likely reflects the outcome of various host and/or symbiont metabolic or cellular pathways which together regulate the expulsion of symbiont cells (coral bleaching) from the host tissue (Weis, 2008). However, initial identification of reef systems or individual coral colonies with desirable traits such as thermal resilience is challenging (Parkinson et al., 2020), often requiring expensive and time-consuming efforts not available to most front-line restoration practitioners. New tools are needed that utilize our collective knowledge of coral physiology and/or genetics to inform on key traits and facilitate colony selection for restoration activities.

The intracellular symbiotic algae (family: Symbiodiniaceae) are typified by high genetic variability within and across individual species (LaJeunesse et al., 2018). In contrast, our understanding of phenotypic variability across these algal species lags behind the genetics, largely due to challenges in measuring cellular characteristics of algae living within the host tissue. Nevertheless, coral thermal tolerance is often tied to specific symbiont species (Abrego et al., 2008; Suggett et al., 2017; van Woesik et al., 2022) and further consideration is needed for how functional traits link to underlying genetic variability across this algal lineage. Bio-optical tools such as the Pulse Amplitude Modulated (PAM) fluorometer that measures algal-specific traits such as variable chlorophyll *a* fluorescence have already provided critical insight into the variability of thermal resilience across coral species, and individual colonies (Warner et al., 1999; Voolstra et al., 2020; Cunning et al., 2021). However, more sophisticated fast repetition rate fluorometers can offer greater insight into algal-centric thermal responses (Hoadley et al., 2019; Hoadley et al., 2021) or functional trait variability (Suggett et al., 2015; Suggett et al., 2022), leading the way towards further integration of these tools for coral research. Recently, we developed a low-cost, multispectral, and fast repetition rate fluorometer capable of generating over 1000 individual metrics within a short (11-minute) timespan and showcased its utility in defining photosynthetic phenotypes across algal genera hosted by seven different coral species under active restoration in the Florida Keys (Hoadley et al., 2023). Importantly, this work and others suggest that algal-centric photo-physiological metrics are correlated with bleaching severity and such information could provide a scalable means for identifying individual colonies, coral species, or reef sites with high tolerance to thermal stress or other desirable traits. However, further exploration is needed to understand if these highly dimensional and algal-centric physiological metrics, along with machine-learning techniques can be effectively utilized to develop predictive models for accurate trait-based selection of reef corals.

Here we evaluate the use of a machine learning technique to generate a predictive model which estimates bleaching tolerance based solely on rapid and non-destructive chlorophyll-a fluorescence, photo-physiological metrics measured with a low-cost and open access bio-optical tool. First, a four-week long thermal bleaching experiment was performed on 156 genotypes (genets) of *Acropora palmata* at a land-based restoration facility in the Florida Keys. Next, correlations between algal photo-physiological metrics and bleaching response (percent change in Fv/Fm or Absorbance at 675nm) were ranked and corals were randomly selected for use in model training or testing. Evaluation was performed using a bootstrap technique to ensure robust model performance across all coral genotypes. Prior studies that have used host genetic information for predicting thermal tolerance found that accuracy improved dramatically when environmental or information on the dominant symbiont type were also incorporated into their model (Fuller et al., 2020). Our study extends this predictive concept by focusing on the underlying phenotype of the symbiont as a tool for assessing coral tolerance. Artificial intelligence-based techniques are increasingly applied within conservation and earth sciences (Evans et al., 2012; Reichstein et al., 2019) and our study demonstrates its utility for trait-based selection of reef corals using low-cost and rapid, bio-optical measurements of symbiont physiology.

## Materials and Methods

### Coral selection and husbandry

Mote’s International Center for Coral Reef Research and Restoration (MML-IC2R3) on Summerland Key, Florida contains approximately 60 land-based raceways, supplied with filtered, UV sterilized, temperature-controlled, near-shore seawater and maintained underneath 60% shade cloth canopies and corrugated clear-plastic rain-guards as needed. Peak midday irradiance within these outdoor raceway aquaria was measured (Walz, 4pi sensor) in 2021 (Hoadley et al., 2023) at ∼400 µmol m^-2^ sec^-1^ under full sunlight, in the month of May. Individual coral genotypes were pulled from Mote’s restoration broodstock, with fragments mounted to ceramic disks (Boston Aqua Farms, 3 cm diameter) using cyanoacrylate gel (Bulk Reef Supply). Of the 156 *A. palmata* within the study, 30 were reared as sexual recruits from a batch cross collected from the Upper Florida Keys in 2013 (29 genets) and 2015 (1 genet), 80 genets were the product of a batch cross sourced from Elbow/Biscayne in 2017, 42 genets were the product of a batch cross sourced from the lower keys in 2020, and 4 were sourced as collections directly from the reef Looe Key, Sand Key, WDR, Turtle reef) in 2018, 2021, 2021, and 2014 respectively. Genotype identification was confirmed using single nucleotide polymorphism loci (Kitchen et al., 2020). All coral fragments utilized in this study had been propagated in Mote’s *ex-situ* nursery and acclimated to the Climate and Acidification Ocean Simulator (CAOS) system for 4 weeks prior to the start of experimental conditions.

### Phenotyping coral photosymbionts using chlorophyll *a* fluorescence

Prior to experimental bleaching, phenotypic measurements were derived (between March 31^st^ – April 5^th^, 2022) from a single coral fragment (ramet) per *A. palmata* genotype. Each fragment was dark acclimated for 20 minutes prior to fluorescence measurements. Importantly, the fragment used for capturing phenotypic data is separate from the two fragments used to assess bleaching response metrics at the end of the experiment. Within-genotype phenotypic variability is thus incorporated into our overall design.

Fluorescence excitation in our phenotyping protocol was achieved using four excitation wavelengths (415, 448, 470 and 505-nm) which preferentially target different photopigments within the symbiotic algae. Fluorescence induction, which consists of the lowest initial fluorescent measurement (F_o_) to maximal fluorescence where the signal appears to plateau (F_m_) consisted of a series of brief excitation pulses, each 1.3-*μ*s long and followed by a 3.4-*μ*s dark interval. For our samples, 32 flashlets was sufficient to reach a plateau in our induction curve which was then used to calculate spectrally dependent Fo and Fm values, along with subsequent metrics ((ΦPSII, NPQ, and qP) as previously described in Hoadley et al., 2023. Additionally, spectrally dependent excitation pressure over PSII (Q_m_) was also calculated using the equation:

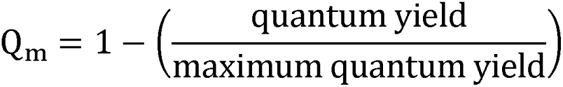

The excitation pressure over PSII equation is adapted from (Iglesias-Prieto et al., 2004) where maximum quantum yield reflects the highest quantum yield value within the actinic light protocol for a given sample and excitation wavelength. Power (irradiance in PAR) estimates were used to determine the spectrally dependent functional absorption cross section of PSII (σ_PSII_) and antennae bed quenching (ABQ) according to previously published methods (Kolber et al., 1998; Oxborough et al., 2012). A 300-millisecond fluorescent relaxation measurement followed immediately after induction and utilized the same 1.3-*μ*s excitation flash but followed by an exponentially increasing dark period (starting with 59-*μs*). This induction and relaxation process was run sequentially for each excitation color with a 200-millisecond delay in between each run. Measured values reflect an average of 6 repeats per sampling time-point. Fluorescent measurements were acquired during a 6-minute actinic light protocol which began with an initial dark period, followed by three different light intensities (200, 50, 400 µmol m^-2^ sec^-1^) and a dark recovery period (See PAR profile in Figure 1). A total of 28 evenly spaced sampling time points were recorded during this actinic light protocol. Definitions and units for each of our 8 spectrally dependent photo-physiological metrics calculated for each sampling time point in our phenotyping protocol can be reviewed in Table 1.

**Figure 1.**
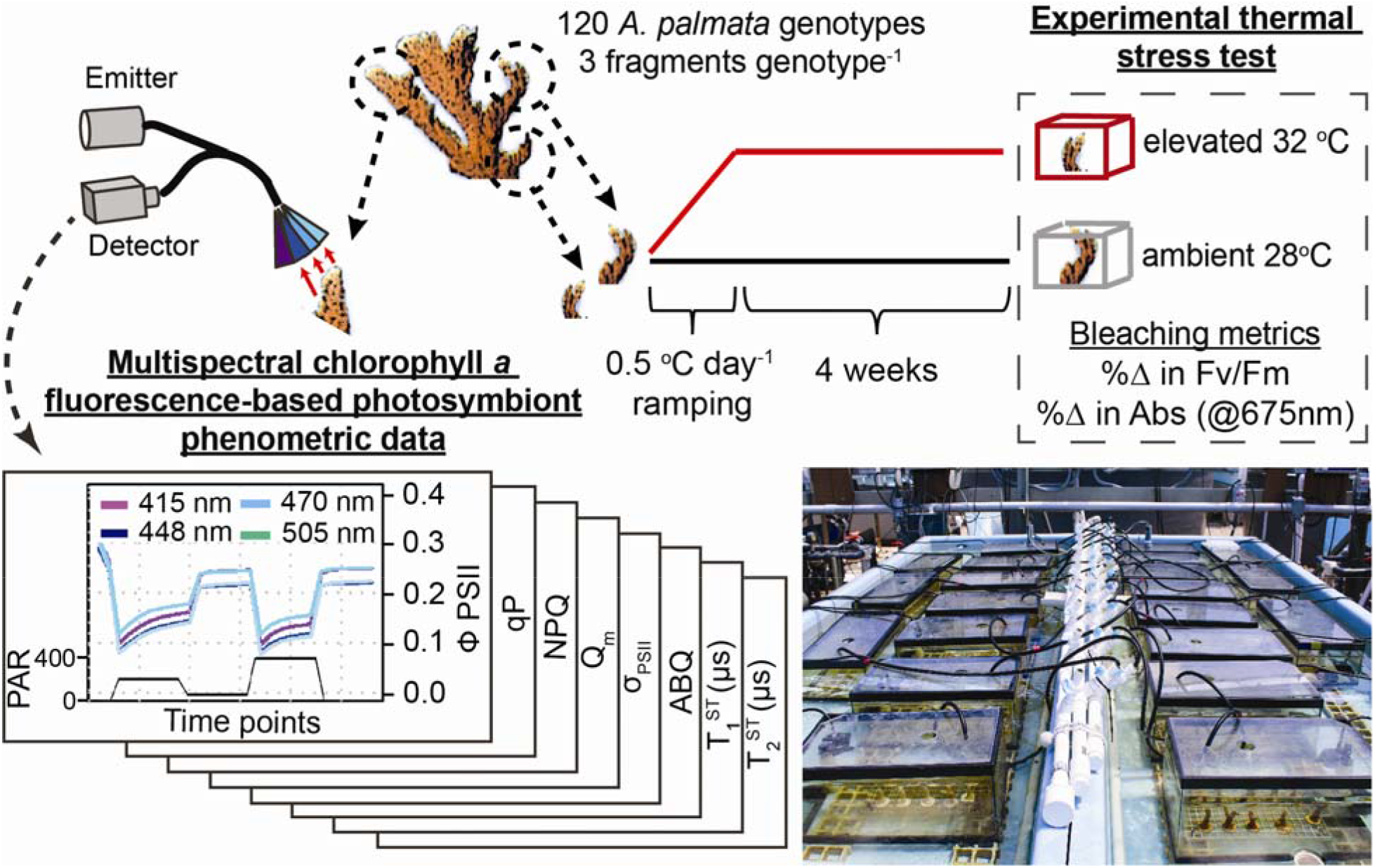
Experimental Setup: For each *Acropora palmata* colony, three fragments were removed for use in the experiment. Chlorophyll-a fluorescence-based measurements were recorded from one fragment using a single turnover, multispectral fluorometer. Bottom left graphs reflect the multispectral data and actinic light protocol used in our analysis. The other two fragments were placed into control and treatment tanks and exposed to a four-week-long thermal experiment. At the end of the thermal experiment, Fv/Fm and Absorbance readings were measured from the two fragments (control and treatment) and utilized to calculate the %Δ in response to high temperature. Bottom right photograph reflects the experimental bleaching raceways and tanks used in the study (corals in photo are *A. cervicornis* and not part of this work).

**Table 1:** Table of photo-physiological parameters: Each defined parameter is represented by spectrally dependent values at each sampling time point (Fig. 1).

| Term | Definition | Units |
| --- | --- | --- |
| $\Phi_{\text{PSII}}$ | Quantum yield of PSII. Measures the proportion of light energy captured by chlorophyll which is then utilized by the PSII reaction center for photosynthesis. | No units |
| qP | Photochemical quenching. Fraction of PSII reaction centers able to utilize light energy for photosynthesis. | No units |
| NPQ | Non-photochemical quenching. Light energy dissipation pathway describing the downregulation of PSII. | No units |
| $\sigma_{\text{PSII}}$ | Absorption cross section of PSII. A measure of photon capture by light harvesting compounds connected to a PSII reaction center | $\text{nm}^2$ |
| ABQ | Antennae Bed Quenching, Light energy dissipation pathway involving reorientation of light-harvesting compounds. | No units |
| $Q_m$ | Excitation pressure over PSII | No units |
| $\tau_1^{\text{ST}}$ | Rate constant for reoxidation of the $Q^a$ site of the D1 protein within the PSII reaction center. | $\mu\text{-seconds}$ |
| $\tau_2^{\text{ST}}$ | Rate constant for reoxidation within and downstream of the plastoquinone pool | $\mu\text{-seconds}$ |

### Thermal bleaching experiment

Using the Climate and Acidification Ocean Simulator (CAOS) system located at MML-IC2R3, a four-week long thermal bleaching experiment was performed on the coral species, *Acropora palmata* (n=156 genotypes) from April 2-May 6, 2022. All coral fragments utilized in this study were acclimated for one month within the CAOS system prior to the start of the experimental treatment. Control and heat treatments consisted of three shallow raceways per treatment however, only a single fragment from one control and one high-temperature raceway was utilized in the present study. Filtered seawater was continuously supplied to each raceway (118 L hr^-1^), with recirculating flow provided by a 5679 L hr^-1^ external pump to supply the heat exchange system. Additional circulation was provided by four submersible pumps (454 L hr^-1^, Dwyer). All experimental systems were maintained underneath 60% shade cloth canopies and clear plastic rain guards with additional 60% shade cloth covers added between 12:00 and 14:00 daily and experienced similar light levels as described above. At the start of the experiment (April 2, 2022), the high-temperature raceway increased from 27.5°C to 31.5°C by 0.5 °C each day (8 days total). Control raceways remained at 27.5°C C for the duration of the 1-month experiment and high-temperature raceways remained at 31.5°C for 3 weeks and 32.0°C for the remainder of the 1-month experiment.

### Bleaching response metrics

Measurements of the maximum quantum yield (Fv/Fm) using 448-nm excitation and absorbance spectra were measured from all experimental fragments on experimental days 39-42 (May 7-8^th^ and 9-10^th^ of 2022 for high temperature and control treatment groups, respectively). Maximum quantum yield measurements were derived from control and high temperature fragments after a 20-minute dark acclimation period and using the same induction and relaxation protocol described above. Absorption-based measurements were calculated on all coral fragments and achieved by measuring the reflectance spectra according to previously established methods (Rodriguez-Román et al., 2006). A custom fiber optic cable (Berkshire Photonics) coupled a white LED (Luxeon) to a USB2000 spectrophotometer (Ocean Optics) for assessing spectral reflectance from all coral fragments in control and treatment conditions. Reflectance measurements were normalized to a bleached *A. palmata* skeleton and then converted into absorbance measurements which can serve as a non-invasive proxy for changes in cell density/chlorophyll *a* content associated with coral bleaching (Rodriguez-Román et al., 2006; Hoadley et al., 2016). Here, absorbance readings were measured at 675nm which reflects the maximum chlorophyll-*a* absorbance band.

### Statistical analysis and bleaching response model generation

All analyses were conducted in R (v.3.5.1) (Team, 2017). For each coral genotype, bleaching response metrics (Fv/Fm and absorbance at 675nm) were calculated as the percent change between control and high temperature fragments (Figure 2). To minimize bias associated with high correlation between certain photophysiological traits, a correlation matrix was first used to identify and then remove individual metrics with high correlation (Rho > 0.99) to one another. All remaining photophysiological metrics were then used to build a phenotypic dendrogram using the R packages pvclust (Suzuki & Shimodaira, 2013) and dendextend (Galili, 2015). Resulting dendrogram with 10,000 bootstrap iterations was then used to cluster individual genotypes into four distinct clusters/phenotypes. Next, significant differences across our cluster-based phenotypes for thermally-induced changes in absorbance and Fv/Fm were measured using a one-way ANOVA with a Tukey-posthoc (all data fit assumptions of normality).

**Figure 2.**
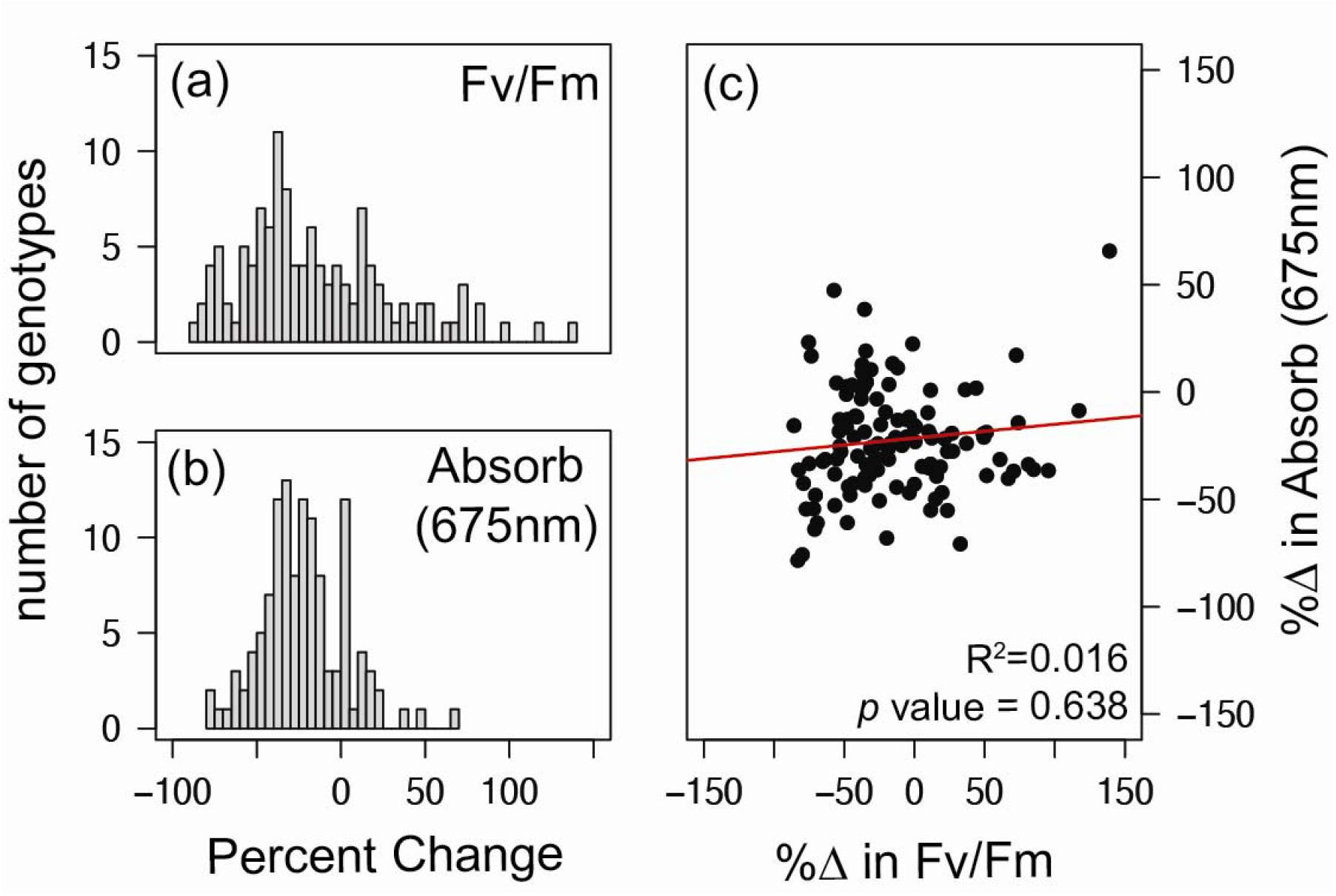
Bleaching Response Metrics: Results of the six-week thermal stress experiment were characterized by recording Fv/Fm and Absorbance (at 675nm) measurements from each coral colony and represented as the percent change between control and high temperature conditions. Distribution of colony responses are displayed as separate histograms for %Δ in Fv/Fm (**a**) and %Δ in Absorbance (**b**). Individual colony responses for both bleaching response metrics are reflected in the correlation plot in panel (**c**) where the red line represents the best linear fit to the data.

For assessing individual photophysiological metrics and their correlation to bleaching response, all photo-physiological parameters were first screened for correlation with one another (autocorrelation scores < 0.85 Rho) and then individually tested for correlation (spearman) with bleaching response metrics (% change in absorbance and Fv/Fm). Only photo-physiological metrics with significant (*p < 0.05*) correlation were included in Figure 4 (and Supplementary tables S1 and S2).

Selecting which photo-physiological metrics to use to optimize a predictive bleaching model is a critical step in our analytical pipeline. First, we screened for autocorrelation between photophysiological metrics. Different correlation cutoff values were used when selecting metrics for the assessment of % change in absorbance (Rho > 0.85) and Fv/Fm (Rho > 0.75) and are based on resulting predictive performance. Next the Boruta R package (Kursa & Rudnicki, 2010) was used to carryout feature selection on the remaining photophysiolical metrics using a random forest based assessment and prioritization. Resulting prioritized photophysiological metrics were then used to develop separate models for predicting percent change in Fv/Fm and absorbance (at 675-nm) between control and treatment fragments. Randomly selected coral genotypes were used in model training which consisted of using the Random Forest (RF) regression algorithm to generate each model iteration. The strength of each individual RF model was then tested by predicting bleaching response on 40 randomly selected colonies which were not used to generate the model. Accuracy of each model was then measured using goodness of fit (R^2^) and the root mean square error (RMSE – describes the model error) between predicted and observed bleaching responses (% change in Fv/Fm or absorbance). In addition to goodness of fit, the 40 ‘test’ corals were also ranked based on model predictions, and then the significance of actual bleaching responses between the top and bottom 10 corals was statistically compared using a t-test (Figure 4c,f). Accuracy or our RF models was evaluated as a function of the number of corals (between 20 and 80) used for training, with each model repeated 100 times with randomly selected corals (Bootstrap resampling technique). Importantly, outcomes were always tested using 40 different corals (also randomly selected and separate from those used in training). Bootstrap model scores enable us to evaluate stability and ensure performance was not biased through inclusion/exclusion of a given coral genotype. Raw data along with analytical scripts for generating Figures 2-5 are available via github (khoadley/bleaching-prediction-2023).

## Results

### Bleaching Response

Of the 156 *A. palmata* colonies that were evaluated for thermal stress resilience, 32 were removed prior to the end of the two-month experiment due to extreme bleaching (> 75%), while another 4 were not properly measured for photophysiology at the onset of the experiment. Because these corals were either pulled from the experiment early, or were missing critical data, they are not included in any downstream analyses or predictive bleaching model testing shown here. For the remaining corals (n=120), the average percent change in bleaching response metrics for Fv/Fm and absorbance was −14.58% (±45% SD) and −23.36% (± 24% SD) respectively (Figure 2a,b). Correlation between the two bleaching response metrics was evaluated using correlation (R^2^ = 0.016, *p* value = 0.639) and reflects no significant linear relationship (Figure 2c).

### Linking photo-physiological phenotypes with bleaching response

Our photophysiology-based dendrogram separated our 120 coral colonies into four distinct clusters with high bootstrap support (Figure 3a). We next wanted to see if significant differences in bleaching sensitivity existed between our four identified clusters/phenotypes. A one-way ANOVA with a Tukey-posthoc found differences in the observed % change in absorbance and Fv/Fm across clusters (*p <* 0.0001). For Absorbance, phenotype 1 displayed a significantly (*p* < 0.044) larger response to thermal stress as compared to phenotypes 2 and 3 (Figure 3b). Phenotypes 2 (*p=*0.01) and 4 (*p=*0.004) also had a significantly larger response to thermal stress as compared to the most resistant phenotype (phenotype3). For high-temperature induced changes in Fv/Fm, phenotype 1 had a significantly (*p <* 0.002) larger reduction as compared to phenotypes 2 and 4 (Figure 3c).

**Figure 3.**
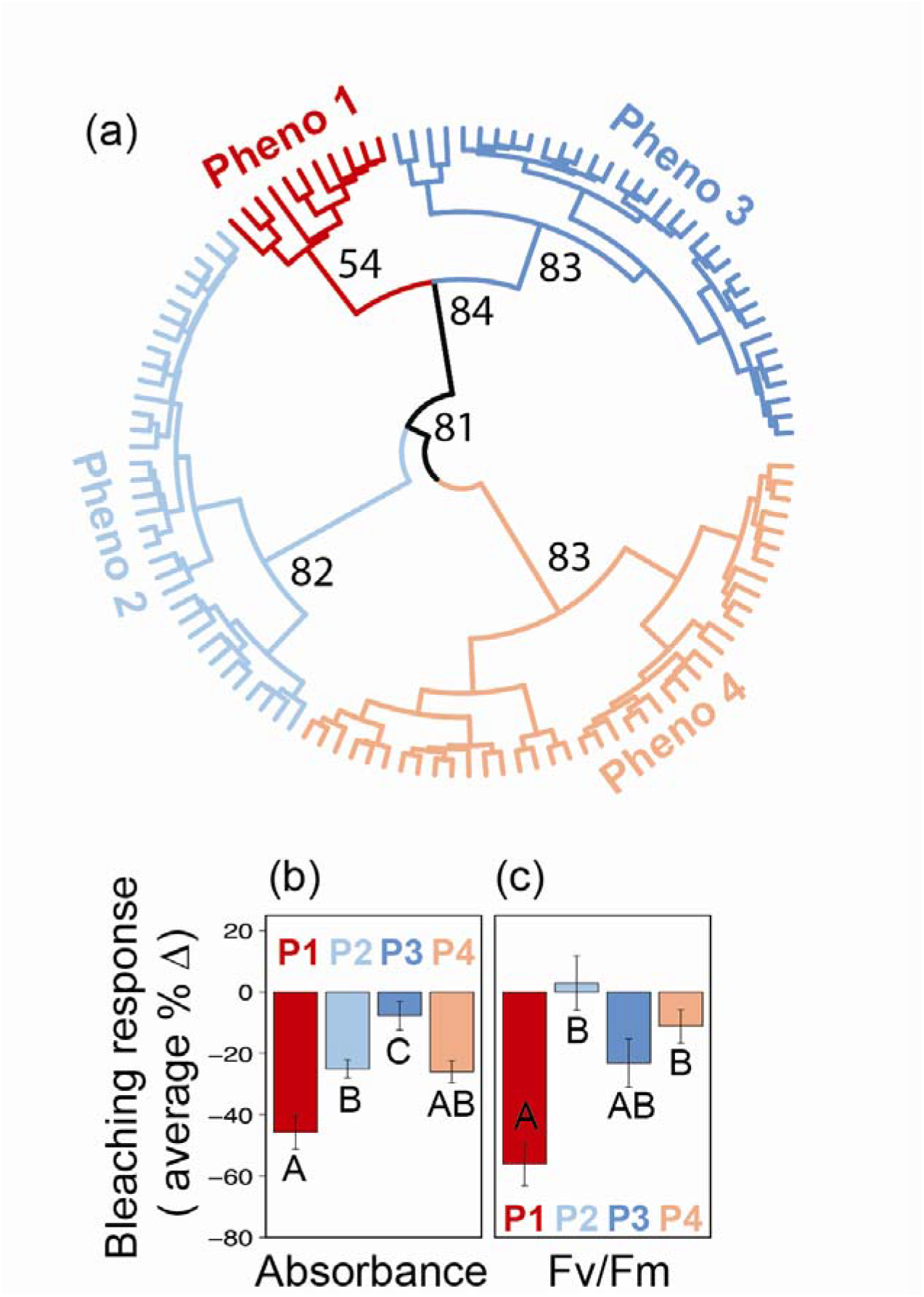
Coral Photosymbiont Phenotypic Variability: Phenomic dendrogram (**a**) derived from 716 photo-physiological metrics (autocorrelation values <0.99 Rho). The largest four clusters are color coded (**a**). Bootstrap values are based on 10,000 iterations and are indicated for major nodes delineating the four phenotypes. Bar graphs reflect the average ± se for observed temperature-induced changes in absorbance (**b**) and Fv/Fm (**c**) for the dendrogram’s four identified clusters/phenotypes. Letters under individual bars represent significant differences across phenotypes as measured using a one-way ANOVA with Tukey-posthoc pairwise comparisons. An extended version of this figure which includes mean traces for each photo-physiological metric can be found in the supplementary document (Fig S1).

### Correlations between algal phenomics and bleaching response metrics

A total of 896 photo-physiological responses (8 photo-physiological responses * 4 excitation wavelengths * 28 sampling time points) were measured as part of our photo-physiologically based phenometric assay. Using data from all 120 colonies, these photo-physiological metrics were first screened for high-correlation (Rho > 0.85) with one another and then those remaining (311 photophysiological metrics) were directly tested for significant correlation (*p* < 0.05) with the two individual bleaching metrics. Only 66 and 170 photo-physiological metrics were deemed to have significant correlation with % change in Absorbance and Fv/Fm respectively. The absolute range in Rho for significant metrics was between 0.179 - 0.448 for Absorbance and 0.184 - 0.455 for Fv/Fm (See Supplemental tables S1 and S2). Correlations were then plotted as a function of which step in the actinic light protocol the phenometric values were derived (Figure 4a,c). For both bleaching response metrics, significant correlations are predominantly made with phenometrics derived immediately (within 10 seconds) after an increase in light intensity (sampling time points 3-5 and 17-19), and strongly suggest these transitional periods contain important photochemical signatures related to how symbionts cope with environmental stress. Next, significant correlations for each bleaching response metric were further evaluated by which photo-physiological metrics they reflect (Figure 4b,d). The most common photo-physiological metrics significantly correlated with %Δ in FvFm were τ_1_^ST^ (28%), τ_2_^ST^ (21%), and antennae bed quenching (37%) while the most common for %Δ in absorbance were the absorbance cross section of PSII (39%), and Antennae Bed Quenching (36%).

**Figure 4.**
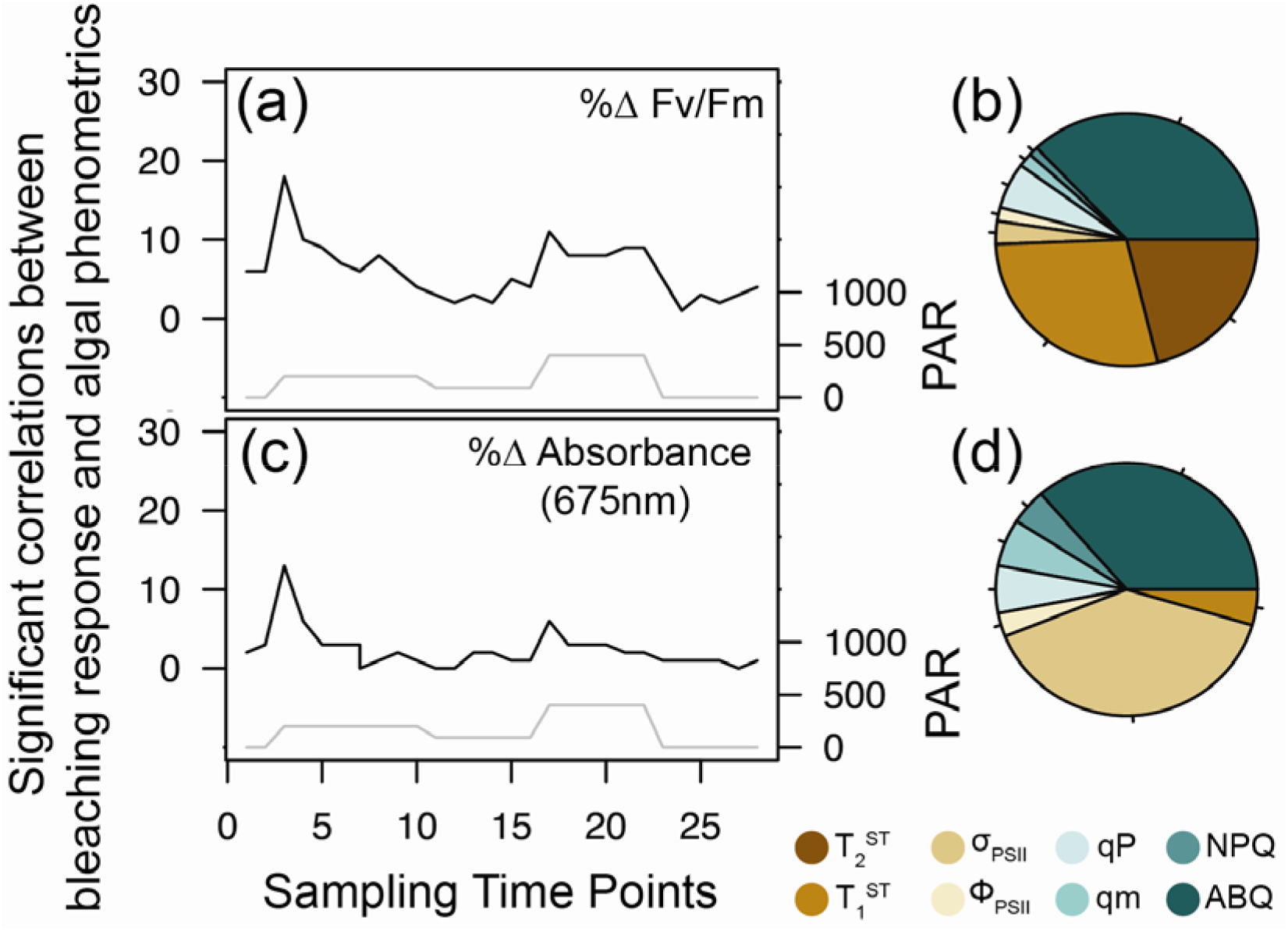
Phenomic vs. Bleaching Response Correlations: A correlation matrix was used to assess relationships between bleaching response metrics (%Δ in Fv/Fm and %Δ in absorbance) and all algal photo-physiological metrics derived from the multispectral single-turnover fluorometer. Panels (**a**) and (**c**) reflect significant correlations (spearman) between photo-physiological metrics and %Δ in Fv/Fm or %Δ in absorbance (170 and 66 total/resulting metrics, respectively) and are summarized according to the sampling time point (black line) from which they are derived. The gray line in each figure reflects the PAR values for each step in the actinic light protocol. Pie charts in panels (**b**) and (**d**) reflect the fraction of each algal metric significantly correlated with %Δ in Fv/Fm and %Δ in absorbance, respectively.

### Predictive bleaching model evaluation

For both bleaching response metrics, Random Forest-based models were able to generate predictions which were significantly correlated with observed trends in thermal bleaching responses (Figure 5b,e). Importantly, average bootstrapped correlations between observed and predicted responses improved by incorporating additional coral genotypes for model generation (Figure 5d,g). However, the bootstrapped-averaged accuracy of predicting changes in Fv/Fm peaked with an R^2^ value of 0.42 (± 0.018 CI, using 80 colonies for model testing). Accuracy in predicting changes in absorbance peaked at 0.33 (± 0.019 CI, using 80 colonies for model testing). While our R^2^ plots suggest that model accuracy may have improved with additional colonies used in training, the root mean squared error for both metrics plateaued at roughly 23 and 19 (for Fv/Fm and absorbance respectively) with at least 60 colonies used for model generation. When used to evaluate the average bleaching response for the top and bottom ranked corals, model results for changes in Fv/Fm were between 67 and 71% accurate when at least 60 colonies were used to generate the model whereas results for changes in absorbance peaked between 89-91% accurate when using at least 60 colonies for model generation (Figure 5c,f).

**Figure 5.**
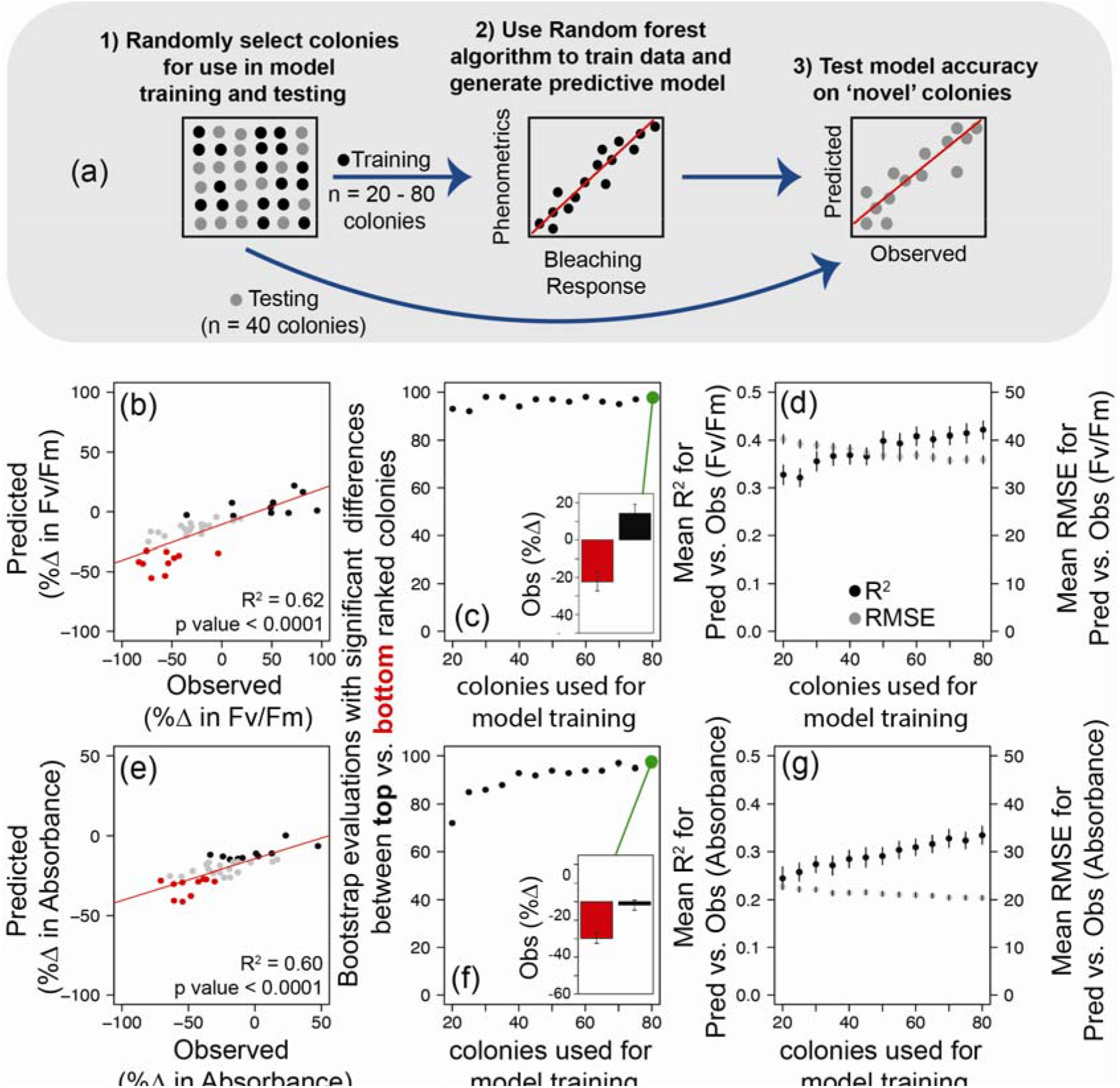
Model Training and Evaluation with Bootstrap Resampling: Overall model performance was evaluated based on the total number of colonies utilized in training (between 20-80 colonies) while model testing was always carried out using a total of 40 colonies. For each evaluation, samples were randomly selected for training and testing as reflected in panel **a** (steps 1-3). Each evaluation process was bootstrapped 100 times to ensure model outcomes were robust and not reflective of any specific colony sample. Representative test data sets reflecting the Predicted vs. Observed bleaching responses as %Δ in Fv/Fm and %Δ in absorbance (675nm) are found in panels (**b**) and (**e**) respectively. Observed bleaching response scores for the top (black) and bottom (red) ranked samples were then tested for significant differences using a t-test. The number (out of 100) of bootstrapped samples with significant (*P* < 0.05) differences in observed bleaching scores between the top and bottom-ranked colonies are reflected in panels **c** (for %Δ in Fv/Fm) and **f** (%Δ in absorbance 675nm). Inset bar graphs in panels **c** and **f** reflect the mean (± standard deviation) observed bleaching responses for the top and bottom-ranked colonies as evaluated with a model using 68 colonies for training. Panels (**d**) and (**g**) reflect the mean (± 95% CI) R^2^ correlation (black dots) and RMSE (gray dots) values between the Observed and Predicted data (as represented in panels **b** and **e)** as a function of the number of colonies utilized to train each model.

## Discussion

Our objective was to determine if easily measured, chlorophyll-a fluorescence-based photo-physiological metrics could be used as a predictive tool for determining thermal tolerance among different genotypes of the coral *Acropora palmata.* Identifying the degree of thermal tolerance among different genotypes of the same coral species (or different corals) is a common goal for many field studies and restoration initiatives (Voolstra et al., 2020; Voolstra et al., 2021; Grummer et al., 2022; Evensen et al., 2023), yet current practices often require labor-intensive experimental procedures, and the outcome is limited to only coral genotypes utilized in the study. Establishing an Artificial Intelligence (AI) based approach where non-invasive measurements can be used to predict thermal tolerance in novel colonies could remove a major bottleneck in trait-based identification/selection of reef corals within basic and applied research settings. Despite low correlation between our two bleaching response metrics (Fv/Fm and absorbance, Fig 2c), our phenomic-based dendrogram indicated that significant differences in bleaching sensitivity existed across identified coral clusters (Figure 3) and suggests that underlying photo-physiological data can be used to forecast thermal response within individual colonies. To test this concept, we trained a Random Forest-based algorithm using algal phenotypic data, along with bleaching response metrics to predict temperature tolerance in novel coral species (Figure 4-5). Such an approach can extend results of an experiment using data derived from bleaching assays to train a model to infer information on novel coral genotypes, thereby vastly increasing the value and utility of an individual experiment. High-throughput assays for identification of thermal stress already rely on photochemical signatures to assess holobiont response (Voolstra et al., 2020; Cunning et al., 2021) and our approach builds on these ideas through a massive increase in the quantity and dimensionality of photosynthetic-data available.

### Phenotypic variability in bleaching response

Bleaching response metrics were evaluated for each coral genotype as a percent change between fragments from control and high temperature treatments. Percent change in Fv/Fm varied broadly across individual genotypes yet collectively averaged close to zero, indicating relative thermal stability of the symbionts (Figure 2a). While *A. palmata* colonies *in situ* are commonly found to host *Symbiodinium fitti* (LaJeunesse, 2002), colonies acclimated to land-based nurseries are often dominated by the more thermally tolerant symbiont, *Durusdinium trenchii* (Gantt et al., 2023) and can retain this symbiosis even two years after transplant back onto nearby reefs (Elder et al., 2023). Indeed, all the colonies used in our experiment were previously housed in Mote’s land-based nursery on Summerland Key (Mote International Center for Coral Reef Research and Restoration) and likely also host the thermally tolerant *D. trenchii* symbiont. Dominance by *D. trenchii* symbionts may explain the minimal average reduction in Fv/Fm observed in response to 30 days of elevated temperatures. However, high variability across genotypes (Figure 2a) may also suggest variability in symbiont dominance across corals, possibly with more *S. fitti* dominated corals removed early in the experiment. In contrast, changes in absorbance were more pronounced, with an average reduction of 23 percent. Reductions in absorbance indicate a decline in photo pigmentation most likely associated with a decline in symbiont cell density or chlorophyll *a* cell^-1^. Despite discordant responses to thermal stress, significant differences in the two bleaching response metrics were still found across cluster-assigned phenotypes (Figure 3). These findings suggest a strong linkage between the underlying photo-physiological poise of the coral colony and its resilience to future thermal stress. Notably, such linkage extends past intra-colony variability as different ramets were used for bleaching response (post-experiment) and photo-physiological assessment (pre-experiment), highlighting the strength and potential utility of our technique within restoration/conservation settings.

Concordant responses in Fv/Fm and absorbance found in prior studies assume that bleaching is the result of thermal damage to the algal photosynthetic apparatus which induces the over-production of radical oxygen species (ROS) which leach out into the host environment and promote the signaling cascade that triggers cell expulsion (bleaching) (Weis, 2008; Hawkins & Davy, 2012; Hawkins et al., 2015). Indeed, this thermal stress pathway is common among many coral species, especially those considered thermally sensitive (Weis et al., 2008; Weis, 2008). However, thermal stress may also reduce ROS scavenging activity by the host (Baird et al., 2009), potentially leading to cell expulsion without any degradation to the photosynthetic apparatus within the symbiont. Such discrepancies can lead to discordant interpretations of bleaching tolerance based on what bleaching response metrics are utilized. Identifying what physiological traits link to these separate responses is thus a critical step towards understanding and predicting discordant patterns of thermal response, such as those observed here.

### Linkage between bleaching response and fluorescent signatures of photosynthetic poise

Photosynthesis to irradiance curves incrementally raise light intensity and monitor photochemical response to understand light stress and acclimation mechanisms in marine algae (Warner et al., 2010). Typically, each incremental light step is followed by a brief (20 second to 5 minute) period to allow for acclimation prior to measuring the photochemical response. However, our protocol continuously records photochemical responses throughout our actinic light protocol, capturing the acclimation phase as well (see Supplemental Figure S1). This unique protocol thus allows for a higher-resolution understanding of rapid photochemical changes in response to variable light. Notably, measurement periods immediately after transitioning from dark (or low light) to higher light intensity were most correlated with bleaching response metrics (Figure 4a,c). Prior studies have highlighted the importance of understanding photochemical responses to quick transitions in light intensity (Allahverdiyeva & Suorsa, 2015; Andersson et al., 2019) and variability is indeed notable across species of Symbiodiniaceae (Hoadley et al., 2023). Better performance via smaller or faster acclimatization to rapid changes in light may serve as a proxy for overall stress mitigation and may explain why these metrics are more correlated with bleaching response as compared with subsequent measurements of each light acclimation step. However, further research that focuses on understanding the phenotypic variability within individual coral species (see Supplemental Figure S1) or even across multiple coral and symbiont types will be required to understand if such connections are prevalent across the Symbiodiniaceae family or if the relative importance of individual photophysiological metrics or acclimatization periods differ across species or environments.

Correlation matrices identified separate algal phenometrics as providing the best correlations with the two bleaching response metrics (Fv/Fm and absorbance, Figure 4b-d). Changes in reoxidation kinetics (τ_1_^ST^ and τ_2_^ST^) represented the largest portion of metrics with a significant correlation to temperature-induced changes in Fv/Fm (Figure 4b) and describe the rate of electron transport through and the downstream of the PSII reaction center (Schuback et al., 2021). Reoxidation kinetics are often used to identify stress or degradation at different sites that may lower or inhibit photosynthetic capacity (Hoadley et al., 2021; Suggett et al., 2022; Hoadley et al., 2023) and may explain the relationship with changes in Fv/Fm observed here (Supplemental Table S2). In contrast, changes in absorbance due to high temperatures were most linked to measurements of the absorption cross-section of PSII which reflects the size of light harvesting compounds connected to a PSII reaction center. Given this direct connection to photopigments, it is perhaps not surprising that this metric was well correlated with the percent change in absorbance during thermal stress (Figure 4d, Supplemental Table S2). Additionally, antennae bed quenching (ABQ) was also well correlated with both bleaching response metrics (absorbance and Fv/Fm) and describes the reorientation of light harvesting pigments to dissipate excess light energy. Changes in ABQ may therefor serve as an important proxy for the bleaching response pathway which starts in the symbiont and leads to host driven cell expulsion. Understanding what traits are best utilized to assess thermal response is critical for evaluating the bleaching phenotype and our results here indicate that each response metric has unique sets of biomarkers.

### AI-based predictive models for coral thermal resilience

Model-based predictions using genomic data have previously been applied to evaluating the thermal tolerance of individual colonies of the coral *A. millepora* (Fuller et al., 2020). However, predictive accuracy was low when based solely on host genetics and improved notably once information on environmental conditions and symbiont dominance was included. On a larger spatial scale, survival rates of coral larvae sourced from different locations have been used to predict which reefs along the Great Barrier Reef are most likely to produce thermally tolerant corals (Quigley & van Oppen, 2022). Indeed, AI-based predictive models for coral research and restoration are already in use but lack the high-throughput applicability required for scalable recommendations. Here, our predictive pipeline first uses a correlation matrix to prioritize individual algal photo-physiological metrics which are then fed into a Random Forest AI model and converted into predictions of coral bleaching severity, concordant with experimentally produced observations after long-term (4-weeks) exposure to high temperature stress (Figure 4 and 5). Although overall average model strength peaked at 0.42 and 0.33 R^2^ (for Fv/Fm and absorbance, respectively), the use of additional colonies for initial training would improve accuracy. In addition, our results do not incorporate colonies with the highest thermal sensitivity as those fragments were removed prior to the end of the experiment and we were thus unable to capture their signal. The lack of low tolerance corals within our training dataset may also have impacted the overall strength of our predictive model. Model improvements through better training data, additional colonies or incorporation of additional (easily measured) traits or measurement time points (to capture the thermally sensitive individuals) could help strengthen our approach, providing a robust and broadly applicable technique. Future work will also need to assess prediction accuracy across coral species with and without thermally tolerant symbiont types to assess how discordant responses (Figure 2) impact overall modeling efforts.

While our method largely focuses on the symbiont photophysiology to make predictions, the coral host’s influence on symbiont physiology is well documented (Enriquez et al., 2005; Enríquez et al., 2017; Wangpraseurt et al., 2017; Xiang et al., 2020; Bollati et al., 2022) and can be measured using chlorophyll-*a* fluorescence techniques (Hoadley et al., 2019). In this context, our technique incorporates direct and indirect metrics of physiology from both the symbiont and host, respectively. However, future predictive techniques that also incorporate additional and direct host-centric physiological metrics such as GFP production or ROS scavenging could further improve predictive accuracy and align with known host genomic traits that infer thermal tolerance (Kenkel et al., 2013a; Dixon et al., 2015; Drury & Lirman, 2021; Rose et al., 2021; Quigley & van Oppen, 2022).

## Conclusion

By capturing a more complex fluorescent signature that includes rapid responses to varying light using a low cost and open-source instrument, our approach allows for more nuanced photo-physiological differences to be identified and then applied within our modeling pipeline. Tools that increase our capacity to evaluate traits in a highly scalable and accessible fashion are well suited for use towards ongoing coral reef restoration initiatives. Here, chlorophyll-*a* fluorescence-based algal phenotyping combined with AI predictive models converts our large physiological datasets into actionable products. The relatively low cost of our instrumentation, along with the potential for broad application of trained models, could be highly beneficial as a tool for rapidly selecting coral colonies with desirable traits. Although our focus for this study was thermal tolerance, light, and water quality stress are also common challenges for coral nurseries and outgrowth operations (Vardi et al., 2021; Voolstra et al., 2021) and acclimatization to these stressors is also regulated through a combination of host and symbiont physiology (Hennige et al., 2010; Suggett et al., 2012; Hoadley & Warner, 2017; Xiang et al., 2020). Using a similar model training/testing approach, our phenotyping pipeline could prove useful for informing on light and nutrient acclimatization traits as well. Future versions of our multispectral/actinic light protocol could serve as a highly scalable and standardized means for trait-based selection and comparison of coral/algal phenotypes across reef systems worldwide.

## Acknowledgements

We thank the staff and interns at Mote’s International Center for Coral Reef Research and Restoration on Summerland Key, FL for their assistance and support. Research activities were made possible through the Florida Fish and Wildlife Conservation Commission (SAL-22-2406-SCRP and Florida Keys National Marine Sanctuary (FKNMS-2015-163-A3) permits. The work was funded by the National Science Foundation, grant nos. 2054885 to K.D. Hoadley and the National Ocean and Atmospheric Administration, grant no. NA21NMF4820300 to E.M and C.D.K. The authors declare no competing interests.

## Author Contributions

C.D.K., E.M., and C.N.K. planned and designed the thermal bleaching experiment. K.H developed the predictive model while G.L built the instrument. T.K., and C.N.K. collected fragments, setup and conducted the thermal experiments. A.M., S.L., and S.D. collected all phenotyping and bleaching response data. K.H. analyzed the data and wrote the manuscript. All authors provided feedback on the manuscript. K.H. agrees to serve as the author responsible for contact and ensures communication.

## Data Availability Statement

All data needed to evaluate the conclusions in the paper are present in the paper and/or the Supplementary Materials. Raw data and analytical scripts for Figures 2 - 5 are available via github (khoadley/bleaching-prediction-2023).

## Notes

### Competing Interest Statement

The authors have declared no competing interest.

## References

Abrego, D., Ulstrup, K.E., Willis, B.L., van Oppen, M.J. (2008). Species-specific interactions between algal endosymbionts and coral hosts define their bleaching response to heat and light stress. Proc Biol Sci. 275, 2273–2282.

Allahverdiyeva, Y., Suorsa…, M. (2015). Photoprotection of photosystems in fluctuating light intensities. … of experimental botany.

Andersson, B., Shen, C., Cantrell, M., Dandy…, D.S. (2019). The fluctuating cell-specific light environment and its effects on cyanobacterial physiology. Plant ….

Baird, A.H., Bhagooli, R., Ralph, P.J., Takahashi, S. (2009). Coral bleaching: the role of the host. Trends in Ecology & Evolution. 24, 16–20.

Barshis, D.J., Birkeland, C., Toonen, R.J., Gates, R.D., Stillman, J.H. (2018). High-frequency temperature variability mirrors fixed differences in thermal limits of the massive coral Porites lobata. Journal of Experimental Biology. 221, jeb188581.

Bollati, E., Lyndby, N.H., D’Angelo, C., Kühl, M., Wiedenmann, J., Wangpraseurt, D. (2022). Green fluorescent protein-like pigments optimize the internal light environment in symbiotic reef building corals. Elife. 11, e73521.

Boström-Einarsson, L., Babcock, R.C., Bayraktarov, E., Ceccarelli, D., Cook, N., Ferse, S.C.A., Hancock, B., Harrison, P., Hein, M., Shaver, E. (2020). Coral restoration–A systematic review of current methods, successes, failures and future directions. PloS one. 15, e0226631.

Caruso, C., Hughes, K., Drury, C. (2021). Selecting heat-tolerant corals for proactive reef restoration. Frontiers in Marine Science. 8, 632027.

Cunning, R., Parker, K.E., Johnson-Sapp, K., Karp, R.F., Wen, A.D., Williamson, O.M., Bartels, E., D’Alessandro, M., Gilliam, D.S., Hanson, G. (2021). Census of heat tolerance among Florida’s threatened staghorn corals finds resilient individuals throughout existing nursery populations. Proceedings of the Royal Society B. 288, 20211613.

Dixon, G.B., Davies, S.W., Aglyamova, G.V., Meyer, E., Bay, L.K., Matz, M.V. (2015). Genomic determinants of coral heat tolerance across latitudes. Science. 348, 1460–1462.

Drury, C., Lirman, D. (2021). Genotype by environment interactions in coral bleaching. Proceedings of the Royal Society B.

Elder, H., Million, W.C., Bartels, E., Krediet, C.J., Muller, E.M., Kenkel, C.D. (2023). Long-term maintenance of a heterologous symbiont association in Acropora palmata on natural reefs. The ISME Journal. 17, 486–489.

Enriquez, S., Mendez, E.R., Iglesias-Prieto, R. (2005). Multiple scattering on corals enhances light absorption by symbiotic algae. Limnology and Oceanography. 50, 1025–1032.

Enríquez, S., Méndez, E.R., Hoegh-Guldberg, O., Iglesias-Prieto, R. (2017). Key functional role of the optical properties of coral skeletons in coral ecology and evolution. Proceedings of the Royal Society B: Biological Sciences. 284, 20161667.

Evans, M.R., Norris, K.J., Benton, T.G. (2012). Predictive ecology: systems approaches. Philosophical Transactions of the Royal Society B: Biological Sciences. 367, 163–169.

Evensen, N.R., Parker, K.E., Oliver, T.A., Palumbi, S.R., Logan, C.A., Ryan, J.S., Klepac, C.N., Perna, G., Warner, M.E., Voolstra, C.R. (2023). The Coral Bleaching Automated Stress System (CBASS): A low-cost, portable system for standardized empirical assessments of coral thermal limits. Limnology and Oceanography: Methods.

Fuller, Z.L., Mocellin, V.J.L., Morris, L.A., Cantin…, N. (2020). Population genetics of the coral *Acropora millepora*: Toward genomic prediction of bleaching. Science.

Galili, T. (2015). dendextend: an R package for visualizing, adjusting and comparing trees of hierarchical clustering. Bioinformatics.

Gantt, S.E., Keister, E.F., Manfroy, A.A., Merck, D.E., Fitt, W.K., Muller, E.M., Kemp, D.W. (2023). Wild and nursery-raised corals: comparative physiology of two framework coral species. Coral Reefs. 1–12.

Grummer, J.A., Booker, T.R., Matthey-Doret, R., Nietlisbach, P., Thomaz, A.T., Whitlock, M.C. (2022). The immediate costs and long-term benefits of assisted gene flow in large populations. Conservation Biology. e13911.

Hawkins, T.D., Davy, S.K. (2012). Nitric oxide production and tolerance differ among Symbiodinium types exposed to heat stress. Plant and Cell Physiology. 53, 1889–1898.

Hawkins, T.D., Krueger, T., Wilkinson, S.P., Fisher, P.L., Davy, S.K. (2015). Antioxidant responses to heat and light stress differ with habitat in a common reef coral. Coral Reefs. DOI 10.1007//s00338-015.

Hennige, S.J., Smith, D.J., Walsh, S.-J., McGinley, M.P., Warner, M.E., Suggett, D.J. (2010). Acclimation and adaptation of scleractinian coral communities along environmental gradients within an Indonesian reef system. Journal of Experimental Marine Biology and Ecology. 391, 143–152.

Hoadley, K.D., Lewis, A.M., Wham, D.C., Pettay, D.T., Grasso, C., Smith, R., Kemp, D.W., LaJeunesse, T.C., Warner, M.E. (2019). Host-symbiont combinations dictate the photo-physiological response of reef-building corals to thermal stress. Sci Rep. 9, 9985.

Hoadley, K.D., Warner, M.E. (2017). Use of open source hardware and software platforms to quantify spectrally dependent differences in photochemical efficiency and functional absorption cross section. Front Mar Sci. doi: 10.3389/fmars.2017.00365.

Hoadley, K.D., Lockridge, G., McQuagge, A., Pahl, K.B., Lowry, S., Wong, S., Craig, Z., Petrik, C., Klepac, C., Muller, E.M. (2023). A phenomic modeling approach for using chlorophyll-a fluorescence-based measurements on coral photosymbionts. Frontiers in Marine Science. 10, 1092202.

Hoadley, K.D., Pettay, D.T., Dodge, D., Warner, M.E. (2016). Contrasting physiological plasticity in response to environmental stress within different cnidarians and their respective symbionts. Coral Reefs. 35, 1–14.

Hoadley, K.D., Pettay, D.T., Lewis, A., Wham, D., Grasso, C., Smith, R., Kemp, D.W., LaJeunesse, T., Warner, M.E. (2021). Different functional traits among closely related algal symbionts dictate stress endurance for vital Indo-Pacific reef-building corals. Global Change Biology.

Hughes, T.P., Anderson, K.D., Connolly, S.R., Heron, S.F., Kerry, J.T., Lough, J.M., Baird, A.H., Baum, J.K., Berumen, M.L., Bridge, T.C. (2018). Spatial and temporal patterns of mass bleaching of corals in the Anthropocene. Science. 359, 80–83.

Hughes, T.P., Kerry, J.T., Álvarez-Noriega, M., Álvarez-Romero, J.G., Anderson, K.D., Baird, A.H., Babcock, R.C., Beger, M., Bellwood, D.R., Berkelmans, R. (2017). Global warming and recurrent mass bleaching of corals. Nature. 543, 373.

Iglesias-Prieto, R., Beltran, V.H., LaJeunesse, T.C., Reyes-Bonilla, H., Thome, P.E. (2004). Different algal symbionts explain the vertical distribution of dominant reef corals in the eastern Pacific. Proceedings of the Royal Society of London. Series B: Biological Sciences. 271, 1757–1763.

Kenkel, C.D., Goodbody Gringley, G., Caillaud, D., Davies, S.W., Bartels, E., Matz, M.V. (2013a). Evidence for a host role in thermotolerance divergence between populations of the mustard hill coral (Porites astreoides) from different reef environments. Molecular Ecology. 22, 4335–4348.

Kenkel, C.D., Matz, M.V. (2016). Gene expression plasticity as a mechanism of coral adaptation to a variable environment. Nat Ecol Evol. 1, 14.

Kenkel, C.D., Meyer, E., Matz, M.V. (2013b). Gene expression under chronic heat stress in populations of the mustard hill coral (*Porites astreoides*) from different thermal environments. Molecular Ecology. 22, 4322–4334.

Kitchen, S.A., Von Kuster, G., Kuntz, K.L.V., Reich, H.G., Miller, W., Griffin, S., Fogarty, N.D., Baums, I.B. (2020). STAGdb: a 30K SNP genotyping array and Science Gateway for Acropora corals and their dinoflagellate symbionts. Scientific reports. 10, 12488.

Klepac, C.N., Eaton, K.R., Petrik, C.G., Arick, L.N., Hall, E.R., Muller, E.M. (2023). Symbiont composition and coral genotype determines massive coral species performance under end-of-century climate scenarios.

Kolber, Z., Prasil, O., Falkowski, P. (1998). Measurements of variable chlorophyll fluorescence using fast repetition rate techniques: defining methodology and experimental protocols. Biochimica et Biophysica Acta (BBA) - Bioenergetics. 1367, 88–107.

Kursa, M.B., Rudnicki, W.R. (2010). Feature selection with the Boruta package. Journal of statistical software. 36, 1–13.

LaJeunesse, T. (2002). Community structure and diversity of symbiotic dinoflagellate populations from a Caribbean coral reef. Marine Biology. 141, 387–400.

LaJeunesse, T.C., Parkinson, J.E., Gabrielson, P.W., Jeong, H.J., Reimer, J.D., Voolstra, C.R., Santos, S.R. (2018). Systematic revision of Symbiodiniaceae highlights the antiquity and diversity of coral endosymbionts. Current Biology. 28, 2570–2580. e6.

Oxborough, K., Moore, C.M., Suggett…, D.J. (2012). Direct estimation of functional PSII reaction center concentration and PSII electron flux on a volume basis: a new approach to the analysis of Fast Repetition Rate …. Limnology and ….

Palumbi, S.R., Barshis, D.J., Traylor-Knowles, N., Bay, R.A. (2014). Mechanisms of reef coral resistance to future climate change. Science. 344, 895–898.

Parkinson, J.E., Baker, A.C., Baums, I.B., Davies, S.W., Grottoli, A.G., Kitchen, S.A., Matz, M.V., Miller, M.W., Shantz, A.A., Kenkel, C.D. (2020). Molecular tools for coral reef restoration: beyond biomarker discovery. Conservation Letters. 13, e12687.

Parkinson, J.E., Banaszak, A.T., Altman, N.S., LaJeunesse, T.C., Baums, I.B. (2015). Intraspecific diversity among partners drives functional variation in coral symbioses. Scientific reports. 5, 15667.

Quigley, K.M., van Oppen, M.J.H. (2022). Predictive models for the selection of thermally tolerant corals based on offspring survival. Nature Communications. 13, 1543.

Reichstein, M., Camps-Valls, G., Stevens, B., Jung, M., Denzler, J., Carvalhais, N. (2019). Deep learning and process understanding for data-driven Earth system science. Nature. 566, 195–204.

Rodriguez-Román, A., Hernández-Pech, X., Thomé, P.E., Enriquez, S., Iglesias-Prieto, R. (2006). Photosynthesis and light utilization in the Caribbean coral *Montastraea faveolata* recovering from a bleaching event. Limnology and Oceanography. 51, 2702–2710.

Rose, N.H., Bay, R.A., Morikawa…, M.K. (2021). Genomic analysis of distinct bleaching tolerances among cryptic coral species. … of the Royal ….

Schuback, N., Tortell, P.D., Berman-Frank, I., Campbell, D.A., Ciotti, A., Courtecuisse, E., Erickson, Z.K., Fujiki, T., Halsey, K., Hickman, A.E. (2021). Single-turnover variable chlorophyll fluorescence as a tool for assessing phytoplankton photosynthesis and primary productivity: opportunities, caveats and recommendations. Frontiers in Marine Science. 8, 690607.

Suggett, D.J., Goyen, S., Evenhuis, C., Szabó, M., Pettay, D.T., Warner, M.E., Ralph, P.J. (2015). Functional diversity of photobiological traits within the genus Symbiodinium appears to be governed by the interaction of cell size with cladal designation. New Phytologist. 208, 370–381.

Suggett, D.J., Dong, L.F., Lawson, T., Lawrenz, E., Torres, L., Smith, D.J. (2012). Light availability determines susceptibility of reef building corals to ocean acidification. Coral Reefs. 32, 327–337.

Suggett, D.J., Nitschke, M.R., Hughes…, D.J. (2022). Toward bio-optical phenotyping of reef-forming corals using Light-Induced Fluorescence Transient-Fast Repetition Rate fluorometry. Limnology and ….

Suggett, D.J., Warner, M.E., Leggat, W. (2017). Symbiotic Dinoflagellate Functional Diversity Mediates Coral Survival under Ecological Crisis. Trends in Ecology and Evolution. 32, 735–745.

Suzuki, R., Shimodaira, H. (2013). Hierarchical clustering with P-values via multiscale bootstrap resampling. R package.

Team, R.C. (2017). R: a language and environment for statistical computing. R Foundation for Statistical Computing, Vienna. https://www.R-project.org.

van Woesik, R., Shlesinger, T., Grottoli, A.G., Toonen, R.J., Vega Thurber, R., Warner, M.E., Marie Hulver, A., Chapron, L., McLachlan, R.H., Albright, R. (2022). Coral-bleaching responses to climate change across biological scales. Global Change Biology.

Vardi, T., Hoot, W.C., Levy, J., Shaver, E., Winters, R.S., Banaszak, A.T., Baums, I.B., Chamberland, V.F., Cook, N., Gulko, D. (2021). Six priorities to advance the science and practice of coral reef restoration worldwide. Restoration Ecology. 29, e13498.

Voolstra, C.R., Buitrago-López, C., Perna, G., Cárdenas, A., Hume, B.C.C., Rädecker, N., Barshis, D.J. (2020). Standardized short-term acute heat stress assays resolve historical differences in coral thermotolerance across microhabitat reef sites. Global Change Biology. 26, 4328–4343.

Voolstra, C.R., Suggett, D.J., Peixoto, R.S., Parkinson, J.E., Quigley, K.M., Silveira, C.B., Sweet, M., Muller, E.M., Barshis, D.J., Bourne, D.G. (2021). Extending the natural adaptive capacity of coral holobionts. Nature Reviews Earth & Environment. 2, 747–762.

Wangpraseurt, D., Holm, J.B., Larkum, A.W.D., Pernice, M., Ralph, P.J., Suggett, D.J., Kühl, M. (2017). In vivo microscale measurements of light and photosynthesis during coral bleaching: evidence for the optical feedback loop. Frontiers in microbiology. 8, 59.

Warner, M., Fitt, W., Schmidt, G. (1999). Damage to photosystem II in symbiotic dinoflagellates: a determinant of coral bleaching. Proceedings of the National Academy of Sciences of the United States of America. 96, 8007–8012.

Warner, M.E., Ralph, P.J., Lesser, M.P. (2010). Chlorophyll Fluorescence in Reef Building Corals. In Chlorophyll fluorescence in aquatic sciences: methods and applications, D.J. Suggett, O. Prasil, M. Borowitzka, eds. Springer), pp. 500.

Weis, V.M., Davy, S.K., Hoegh-Guldberg, O., Rodriguez-Lanetty, M., Pringle, J.R. (2008). Cell biology in model systems as the key to understanding corals. Trends in Ecology & Evolution. 23, 369–376.

Weis, V.M. (2008). Cellular mechanisms of Cnidarian bleaching: stress causes the collapse of symbiosis. Journal of Experimental Biology. 211, 3059–3066.

Xiang, T., Lehnert, E., Jinkerson, R.E., Clowez, S., Kim, R.G., DeNofrio, J.C., Pringle, J.R., Grossman, A.R. (2020). Symbiont population control by host-symbiont metabolic interaction in Symbiodiniaceae-cnidarian associations. Nature communications. 11, 1–9.

