## supplemental information for "Bio-optical signatures of *insitu* photosymbionts predict bleaching severity prior to thermal stress in the Caribbean coral species *Acropora palmata*"

**This PDF file includes:**

Extended Phenotypic Variability Figure S1

Correlation Table S1

Correlation Table S2

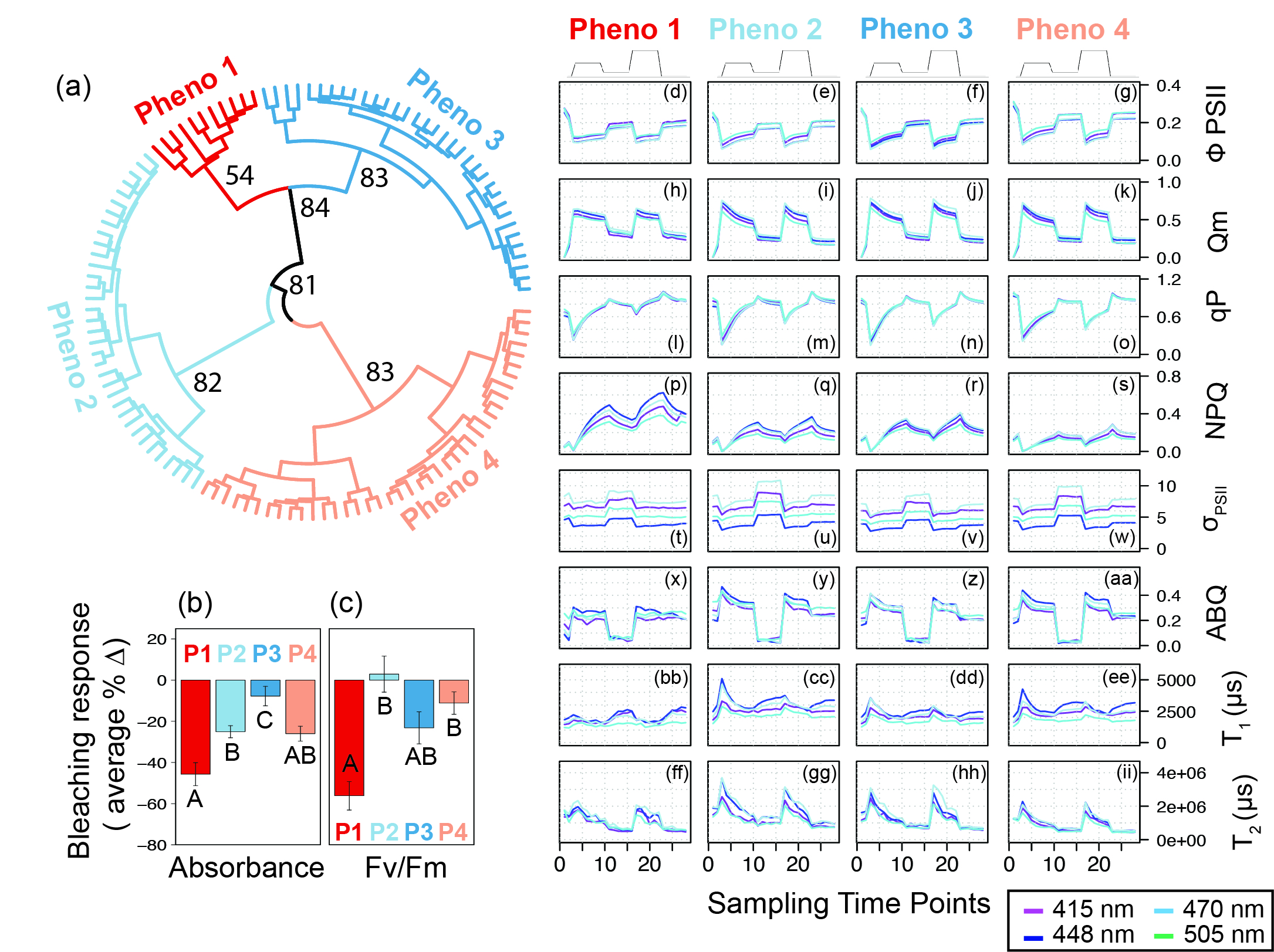

**Figure S1 – Coral Photosymbiont Phenotypic Variability:** Phenomic dendrogram (**a**) derived from 716 photo-physiological metrics (autocorrelation values <0.99 Rho). The largest four clusters are color coded (**a**). Bootstrap values are based on 10,000 iterations and are indicated for major nodes delineating the four phenotypes. Bar graphs reflect the average ± se for observed temperature-induced changes in absorbance (**b**) and Fv/Fm (**c**) for the four identified clusters/phenotypes from the dendrogram. Letters under individual bars represent significant differences across phenotypes as measured using a one-way ANOVA with a Tukey-poshoc. Mean photo-physiological traces for each identified phenotype are reflected in panels d-ii where line color indicates excitation wavelength with purple representing 415-nm; dark blue, 448-nm; light blue, 470-nm; and teal blue, 505-nm. The grey line directly above panels d-g displays the variable light protocol. Panels **d-g** reflect the Quantum Yield of PSII (Φ_PSII_), **h-k** reflects the partial pressure over PSII (Qm), **l-o** reflects photochemical quenching (qP), **p-s** reflects non-photochemical quenching (NPQ), **t-w** reflects the absorption cross-section of PSII **(**σ_PSII_), **x-aa** reflects antennae bed quenching, **bb-ee** and **ff-ii** reflects the reoxidation constants τ_1_^ST^ and τ_2_^ST^ respectively.

**Table S1:** 170 Significant correlations between algal photophysiological metrics and the percent change in Fv/Fm. Correlations performed after screening out photo-physiological metrics with greater than 0.85 Rho correlation with one another.

| Photo-physiological metric | Sampling Time Point | Excitation color (nm) | Spearman Rho | P value |
| --- | --- | --- | --- | --- |
| ABQ | 21 | 505 | 0.45539358 | 1.73E-07 |
| ABQ | 4 | 505 | 0.43836992 | 5.51E-07 |
| Tau1 | 6 | 448 | 0.43713461 | 5.98E-07 |
| ABQ | 19 | 470 | 0.43686121 | 6.09E-07 |
| ABQ | 6 | 415 | 0.41946328 | 1.85E-06 |
| ABQ | 4 | 415 | 0.41791345 | 2.04E-06 |
| ABQ | 19 | 505 | 0.41560417 | 2.35E-06 |
| Tau1 | 5 | 470 | 0.41352858 | 2.67E-06 |
| Tau1 | 5 | 415 | 0.41000784 | 3.31E-06 |
| ABQ | 7 | 415 | 0.40463108 | 4.57E-06 |
| ABQ | 6 | 505 | 0.40272262 | 5.11E-06 |
| ABQ | 20 | 505 | 0.39877539 | 6.45E-06 |
| Tau1 | 6 | 415 | 0.39275184 | 9.12E-06 |
| Tau1 | 3 | 470 | 0.38986796 | 1.07E-05 |
| ABQ | 17 | 505 | 0.38825375 | 1.18E-05 |
| ABQ | 3 | 470 | 0.38664731 | 1.29E-05 |
| Tau1 | 4 | 415 | 0.38167037 | 1.70E-05 |
| ABQ | 3 | 415 | 0.37902351 | 1.96E-05 |
| Tau1 | 8 | 448 | 0.37896735 | 1.97E-05 |
| ABQ | 22 | 505 | 0.37817708 | 2.06E-05 |
| Tau1 | 11 | 415 | 0.37816397 | 2.06E-05 |
| ABQ | 2 | 448 | 0.37426572 | 2.54E-05 |
| Tau1 | 7 | 415 | 0.37196545 | 2.88E-05 |
| ABQ | 18 | 415 | 0.36948775 | 3.28E-05 |
| ABQ | 15 | 448 | -0.3688977 | 3.38E-05 |
| ABQ | 4 | 470 | 0.36718167 | 3.70E-05 |
| Tau1 | 18 | 448 | 0.3643759 | 4.29E-05 |
| ABQ | 9 | 505 | 0.36064712 | 5.20E-05 |
| Tau1 | 8 | 415 | 0.35816284 | 5.91E-05 |
| Tau1 | 8 | 470 | 0.3566423 | 6.38E-05 |
| Tau1 | 18 | 470 | 0.35651903 | 6.42E-05 |
| ABQ | 5 | 470 | 0.35577054 | 6.67E-05 |
| Tau1 | 3 | 448 | 0.35366891 | 7.42E-05 |
| Tau1 | 17 | 505 | 0.35341385 | 7.51E-05 |
| ABQ | 3 | 505 | 0.35286513 | 7.72E-05 |
| ABQ | 18 | 470 | 0.35170962 | 8.18E-05 |
| ABQ | 3 | 448 | 0.35153628 | 8.25E-05 |
| Tau1 | 5 | 448 | 0.35002554 | 8.89E-05 |
| ABQ | 21 | 470 | 0.34938718 | 9.18E-05 |
| ABQ | 5 | 415 | 0.34588407 | 0.00010908 |
| Tau1 | 6 | 505 | 0.34512255 | 0.00011322 |
| Tau1 | 3 | 505 | 0.34055625 | 0.00014125 |
| ABQ | 17 | 415 | 0.3377371 | 0.00016164 |
| ABQ | 6 | 448 | 0.33046699 | 0.00022752 |
| ABQ | 7 | 470 | 0.32711015 | 0.00026567 |
| Tau1 | 19 | 470 | 0.32674475 | 0.00027017 |
| ABQ | 2 | 505 | 0.32486774 | 0.00029437 |
| ABQ | 21 | 415 | 0.32270535 | 0.00032473 |
| Tau1 | 21 | 470 | 0.31954784 | 0.00037429 |
| Tau2 | 18 | 448 | 0.31905522 | 0.00038262 |
| qm | 3 | 415 | 0.31767841 | 0.00040683 |
| ABQ | 1 | 448 | 0.31648709 | 0.00042891 |
| Tau1 | 23 | 415 | 0.31616782 | 0.00043501 |
| Tau1 | 2 | 505 | 0.31510062 | 0.00045599 |
| Tau1 | 2 | 415 | 0.3145127 | 0.00046794 |
| Tau1 | 7 | 505 | 0.31205716 | 0.00052104 |
| ABQ | 10 | 415 | 0.30850303 | 0.00060777 |
| Tau1 | 1 | 448 | 0.30839839 | 0.00061052 |
| qP | 25 | 470 | -0.3067315 | 0.00065578 |
| Tau2 | 6 | 448 | 0.30558996 | 0.00068854 |
| Tau1 | 21 | 415 | 0.30194757 | 0.00080336 |
| Quant | 17 | 448 | -0.3002347 | 0.0008632 |
| ABQ | 22 | 470 | 0.29996662 | 0.00087293 |
| Tau2 | 12 | 470 | 0.29955265 | 0.00088814 |
| ABQ | 2 | 470 | 0.29697649 | 0.00098842 |
| ABQ | 17 | 448 | 0.29651274 | 0.00100754 |
| Tau1 | 1 | 415 | 0.29534122 | 0.00105734 |
| qP | 23 | 448 | -0.293528 | 0.00113886 |
| Tau1 | 4 | 505 | 0.28948526 | 0.00134167 |
| Tau2 | 5 | 415 | 0.28821229 | 0.00141203 |
| Tau1 | 4 | 448 | 0.2853804 | 0.00158074 |
| ABQ | 4 | 448 | 0.28472578 | 0.00162227 |
| Tau1 | 5 | 505 | 0.28351443 | 0.00170172 |
| Tau1 | 25 | 470 | 0.28283444 | 0.00174785 |
| qP | 17 | 415 | -0.2826497 | 0.00176058 |
| Tau2 | 18 | 470 | 0.28256073 | 0.00176674 |
| Tau2 | 26 | 448 | 0.28184123 | 0.00181728 |
| Tau1 | 21 | 448 | 0.28031221 | 0.00192906 |
| ABQ | 22 | 415 | 0.27994418 | 0.00195688 |
| Tau2 | 7 | 470 | 0.27806953 | 0.00210431 |
| Tau1 | 10 | 415 | 0.2776377 | 0.00213966 |
| Tau1 | 9 | 415 | 0.27636382 | 0.00224711 |
| ABQ | 5 | 448 | 0.27474899 | 0.00239033 |
| ABQ | 8 | 415 | 0.27253909 | 0.00259968 |
| ABQ | 20 | 415 | 0.27229085 | 0.0026242 |
| qP | 22 | 415 | -0.2716993 | 0.00268349 |
| Quant | 17 | 415 | -0.2693797 | 0.00292777 |
| qP | 1 | 505 | 0.26889689 | 0.00298106 |
| qP | 3 | 470 | -0.2658823 | 0.00333406 |
| Tau1 | 20 | 415 | 0.26428703 | 0.00353568 |
| Tau2 | 24 | 448 | 0.26340783 | 0.00365143 |
| Sigma | 3 | 415 | -0.2617935 | 0.00387283 |
| Tau1 | 19 | 415 | 0.2617355 | 0.003881 |
| ABQ | 8 | 470 | 0.26078747 | 0.00401682 |
| ABQ | 20 | 470 | 0.26030864 | 0.00408703 |
| Tau1 | 20 | 505 | 0.25814541 | 0.00441809 |
| Tau2 | 16 | 470 | 0.25791557 | 0.00445464 |
| Tau1 | 22 | 505 | 0.25585276 | 0.00479496 |
| ABQ | 2 | 415 | 0.25544364 | 0.00486515 |
| Tau2 | 15 | 470 | 0.25309255 | 0.00528661 |
| Tau1 | 28 | 505 | 0.25296582 | 0.00531023 |
| Tau2 | 15 | 448 | 0.25257692 | 0.00538329 |
| ABQ | 19 | 415 | 0.25168383 | 0.00555446 |
| ABQ | 9 | 448 | 0.25054806 | 0.00577915 |
| ABQ | 9 | 415 | 0.24917787 | 0.00606095 |
| ABQ | 25 | 415 | 0.24782281 | 0.00635159 |
| Tau2 | 27 | 448 | 0.2453794 | 0.0069071 |
| ABQ | 7 | 448 | 0.24528707 | 0.00692891 |
| Tau1 | 17 | 415 | 0.24148199 | 0.00788259 |
| Tau2 | 3 | 448 | 0.24115399 | 0.00796999 |
| NPQ | 4 | 505 | -0.2396989 | 0.00836814 |
| ABQ | 1 | 415 | 0.23891855 | 0.00858882 |
| ABQ | 10 | 470 | 0.23886984 | 0.00860276 |
| Tau2 | 17 | 505 | 0.23884345 | 0.00861033 |
| Tau2 | 13 | 470 | 0.23780649 | 0.00891218 |
| Tau2 | 5 | 470 | 0.23753069 | 0.00899401 |
| ABQ | 18 | 448 | 0.23750626 | 0.00900129 |
| Tau2 | 16 | 448 | 0.23693186 | 0.00917399 |
| Sigma | 3 | 448 | -0.2354576 | 0.00963064 |
| Tau2 | 28 | 415 | 0.23258238 | 0.01057914 |
| NPQ | 3 | 415 | -0.2320653 | 0.01075815 |
| Tau2 | 23 | 448 | 0.23170259 | 0.01088531 |
| Tau2 | 11 | 470 | 0.23138889 | 0.01099634 |
| qm | 3 | 505 | 0.23037337 | 0.01136262 |
| Tau2 | 20 | 505 | 0.23028246 | 0.01139593 |
| ABQ | 9 | 470 | 0.22918998 | 0.01180291 |
| Tau1 | 18 | 415 | 0.22845305 | 0.01208457 |
| ABQ | 8 | 448 | 0.22826387 | 0.01215782 |
| qP | 4 | 415 | -0.2273805 | 0.012505 |
| Tau2 | 28 | 448 | 0.22405741 | 0.01388993 |
| Tau1 | 19 | 505 | 0.22306184 | 0.01433005 |
| Tau1 | 1 | 505 | 0.2199415 | 0.01578909 |
| ABQ | 28 | 415 | 0.21966128 | 0.01592622 |
| Tau1 | 22 | 415 | 0.21965767 | 0.015928 |
| ABQ | 14 | 448 | -0.2152406 | 0.01823035 |
| qP | 3 | 415 | -0.2152089 | 0.01824787 |
| Sigma | 14 | 505 | 0.21459072 | 0.01859228 |
| Quant | 22 | 448 | -0.2141019 | 0.01886858 |
| Tau1 | 21 | 505 | 0.21362801 | 0.01913983 |
| Tau1 | 9 | 505 | 0.21327123 | 0.01934625 |
| Tau2 | 16 | 415 | 0.21326527 | 0.01934971 |
| Tau2 | 26 | 415 | 0.21276575 | 0.019642 |
| Tau1 | 27 | 415 | 0.21157522 | 0.020354 |
| ABQ | 23 | 505 | 0.20907125 | 0.02192433 |
| ABQ | 19 | 448 | 0.20902858 | 0.02195197 |
| Tau2 | 20 | 415 | 0.20845406 | 0.02232701 |
| qm | 17 | 415 | 0.20406835 | 0.02537541 |
| qP | 15 | 470 | -0.2020156 | 0.02692018 |
| ABQ | 16 | 415 | 0.20195674 | 0.02696567 |
| Sigma | 4 | 415 | -0.2018972 | 0.0270117 |
| ABQ | 23 | 415 | 0.20185745 | 0.02704249 |
| Tau2 | 3 | 505 | 0.20145199 | 0.02735813 |
| Tau2 | 15 | 415 | 0.20034438 | 0.02823643 |
| Tau2 | 8 | 448 | 0.19974511 | 0.02872155 |
| Tau1 | 22 | 448 | 0.19847646 | 0.02977195 |
| Tau2 | 11 | 448 | 0.19816503 | 0.03003474 |
| qP | 3 | 448 | -0.1979541 | 0.03021385 |
| Tau2 | 13 | 415 | 0.1958718 | 0.03203102 |
| ABQ | 21 | 448 | 0.19392111 | 0.03381642 |
| Tau2 | 8 | 470 | 0.19347165 | 0.03423949 |
| Sigma | 3 | 470 | -0.1932948 | 0.0344072 |
| Tau2 | 21 | 415 | 0.1917103 | 0.03594074 |
| Tau2 | 19 | 470 | 0.19047448 | 0.0371764 |
| ABQ | 27 | 415 | 0.19024544 | 0.03740929 |
| Tau2 | 13 | 448 | 0.1901098 | 0.03754779 |
| Tau2 | 20 | 448 | 0.19006151 | 0.0375972 |
| ABQ | 10 | 448 | 0.18928034 | 0.03840412 |
| ABQ | 12 | 415 | -0.1872593 | 0.04055947 |
| ABQ | 22 | 448 | 0.18538961 | 0.04264266 |
| Tau2 | 17 | 415 | 0.18367927 | 0.0446258 |

**Table S2:** 66 significant correlations between algal photophysiological metrics and the percent change in Absorbance at 675nm. Correlations performed after screening out photo-physiological metrics with greater than 0.85 Rho correlation with one another.

| Photo-physiological metric | Sampling Time Point | Excitation color (nm) | Spearman Rho | P value |
| --- | --- | --- | --- | --- |
| qm | 3 | 415 | 0.44850336 | 2.79E-07 |
| Sigma | 3 | 470 | -0.4420395 | 4.32E-07 |
| qm | 3 | 505 | 0.43738966 | 5.88E-07 |
| qP | 3 | 470 | -0.4335274 | 7.57E-07 |
| qP | 3 | 415 | -0.4240694 | 1.39E-06 |
| Sigma | 3 | 415 | -0.4016759 | 5.44E-06 |
| qP | 4 | 415 | -0.394328 | 8.34E-06 |
| Sigma | 3 | 448 | -0.3894091 | 1.10E-05 |
| ABQ | 3 | 470 | 0.37965367 | 1.90E-05 |
| Sigma | 4 | 448 | -0.3762535 | 2.28E-05 |
| qP | 3 | 448 | -0.3723145 | 2.82E-05 |
| Sigma | 5 | 448 | -0.3607959 | 5.16E-05 |
| Sigma | 3 | 505 | -0.3530715 | 7.64E-05 |
| Quant | 17 | 415 | -0.3445667 | 0.00011633 |
| Sigma | 17 | 448 | -0.3426989 | 0.00012738 |
| qm | 17 | 415 | 0.33910241 | 0.00015144 |
| Sigma | 4 | 415 | -0.3310656 | 0.00022128 |
| Sigma | 6 | 448 | -0.3298689 | 0.00023392 |
| NPQ | 3 | 415 | -0.3221303 | 0.00033328 |
| Sigma | 18 | 448 | -0.3147719 | 0.00046264 |
| Sigma | 6 | 415 | -0.3122137 | 0.0005175 |
| ABQ | 7 | 470 | 0.30957022 | 0.00058043 |
| Sigma | 5 | 415 | -0.3036699 | 0.00074704 |
| ABQ | 14 | 448 | -0.2923651 | 0.00119412 |
| Sigma | 19 | 448 | -0.2906332 | 0.00128097 |
| Sigma | 1 | 448 | -0.2830845 | 0.00173076 |
| ABQ | 2 | 470 | 0.28078163 | 0.0018941 |
| Sigma | 17 | 415 | -0.2753293 | 0.00233794 |
| Quant | 17 | 448 | -0.275241 | 0.00234584 |
| Sigma | 19 | 415 | -0.2747372 | 0.0023914 |
| Sigma | 7 | 415 | -0.2708647 | 0.00276917 |
| ABQ | 19 | 470 | 0.26573407 | 0.00335235 |
| ABQ | 20 | 470 | 0.2627306 | 0.00374289 |
| ABQ | 5 | 470 | 0.26030298 | 0.00408786 |
| Sigma | 20 | 448 | -0.2599923 | 0.00413401 |
| Sigma | 22 | 448 | -0.2598471 | 0.00415573 |
| Sigma | 1 | 415 | -0.258706 | 0.00433007 |
| Sigma | 18 | 415 | -0.2566446 | 0.00466167 |
| NPQ | 2 | 448 | 0.25476182 | 0.00498418 |
| ABQ | 18 | 470 | 0.25437643 | 0.0050526 |
| Sigma | 7 | 448 | -0.2522978 | 0.00543627 |
| ABQ | 13 | 448 | -0.2508079 | 0.00572704 |
| ABQ | 25 | 448 | -0.2505644 | 0.00577586 |
| ABQ | 4 | 470 | 0.24410994 | 0.00721238 |
| Sigma | 21 | 448 | -0.243569 | 0.00734606 |
| Tau2 | 6 | 415 | 0.24162574 | 0.00784455 |
| ABQ | 15 | 505 | -0.2353846 | 0.00965377 |
| ABQ | 14 | 505 | -0.2301864 | 0.01143121 |
| ABQ | 23 | 505 | -0.2260575 | 0.01304122 |
| ABQ | 2 | 448 | 0.22329389 | 0.01422639 |
| ABQ | 16 | 448 | -0.2178741 | 0.01682537 |
| ABQ | 9 | 470 | 0.21737492 | 0.01708423 |
| Sigma | 17 | 505 | -0.2167067 | 0.01743618 |
| Sigma | 20 | 415 | -0.2127908 | 0.01962728 |
| ABQ | 28 | 470 | -0.2080614 | 0.02258649 |
| ABQ | 26 | 448 | -0.2059448 | 0.02403016 |
| ABQ | 3 | 448 | 0.20025812 | 0.02830582 |
| ABQ | 3 | 415 | 0.1993773 | 0.02902279 |
| Sigma | 10 | 415 | -0.1990786 | 0.02926941 |
| ABQ | 13 | 470 | -0.1972396 | 0.03082726 |
| qm | 9 | 415 | 0.1936143 | 0.03410474 |
| Tau2 | 21 | 415 | 0.18794102 | 0.0398214 |
| ABQ | 22 | 470 | 0.18562202 | 0.04237894 |
| NPQ | 4 | 505 | -0.1823287 | 0.04624559 |
| Tau2 | 4 | 505 | 0.18099857 | 0.04788824 |
| ABQ | 24 | 415 | -0.1799989 | 0.04915449 |
